# Biop-C: A Method for Chromatin Interactome Analysis of Solid Cancer Needle Biopsy Samples

**DOI:** 10.1101/2021.01.11.426176

**Authors:** Sambhavi Animesh, Ruchi Choudhary, Xin Yi Ng, Joshua Kai Xun Tay, Wan-Qin Chong, Boon Cher Goh, Melissa Jane Fullwood

## Abstract

A major challenge in understanding the 3D genome organization of cancer samples is the lack of a method adapted to solid cancer needle biopsy samples. Here we developed Biop-C, a modified *in situ* Hi-C method, and applied it to characterize three nasopharyngeal cancer patient samples. We identified Topologically-Associated Domains (TADs), chromatin interaction loops, and Frequently Interacting regions (FIREs) at key oncogenes in nasopharyngeal cancer from Biop-C heat maps. Our results demonstrate the utility of our Biop-C method in investigating the 3D genome organization in solid cancers, and the importance of 3D genome organization in regulating oncogenes in nasopharyngeal cancer.

## Background

The three-dimensional (3D) genome organization of the nucleus plays a vital role in the regulation of transcription [1,2]. Alterations in 3D genome organization structures including Topologically-Associated Domain boundaries and chromatin loops have been shown to lead to oncogene expression and cancer progression [1,3–6].

High-throughput chromosome conformation capture technologies such as Hi-C have been used to investigate three-dimensional (3D) chromatin conformation [7,8]. The standard Hi-C approach generally requires approximately 1 million cells. Consequently, most previous analyses of cancer samples have been restricted to human cancer cell lines [9–11], but recently, Hi-C has been conducted on clinical samples from liquid cancers such as T-cell Acute Lymphoblastic Leukaemia (T-ALL) [12] and Diffuse Large B-cell Lymphoma (DLBCL) [13] and one solid cancer - gastric cancer [14]. For these cancers, it is possible to obtain 1 million cells – for example, gastric cancers can grow to a large size. However, there are many cancers for which only needle biopsies are available [15].

To allow interrogation of samples with more limited quantities of starting materials, there have been several efforts to reduce the number of cells required to just 1K or 500 cells using modified protocols such as small-scale *in situ* Hi-C (sisHi-C) [16] and Low-C [13]. However, when dealing with solid cancers, a second challenge is that the tissue requires special preparation in order to dissociate the tissue into single cells for Hi-C analysis. The core needle biopsies pose the challenge of both limited cell numbers as well as the requirement for tissue dissociation, which might lead to further loss or degradation of chromatin for analysis. Solid cancers represent approximately 90% of adult human cancers [17], therefore, an easy-to-use method for preparing Hi-C libraries from needle biopsy cancer samples would advance our understanding of how chromatin organization contributes to cancer pathogenesis in solid cancers.

Here, we present Biop-C, a modified *in situ* Hi-C method for the chromatin analysis in solid cancer tissues from needle biopsy samples. The Biop-C method has been designed to be used on small tissue samples obtained from needle biopsies. To demonstrate the utility of this method, we analyzed three Nasopharyngeal Cancer (NPC) patient samples. NPC is an epithelial malignancy of the nasopharyngeal mucosa, and is an aggressive subtype of head and neck cancer [18,19,20]. NPCs are further subdivided into three subtypes *viz* Non-keratinizing undifferentiated carcinoma, Non-keratinizing differentiated carcinoma and Keratinizing squamous cell carcinoma. Depending on the treatment given, the stage of the cancer, and the site where the cancer presents at, NPCs can be small and analysed using core needle biopsies.

It has been established that NPC has a comparatively low mutational burden, and oncogenicity is driven by epigenetic regulation. Typically, NPC associated with Epstein-Barr Virus (EBV) are characterized as having comparatively low DNA mutation rates but widespread DNA hypermethylation and overexpression or mutation of DNA methylation enzymes, histone modification enzymes and chromatin remodelling enzymes [16,17]. Hence, unravelling the 3D conformational structure will provide further insight into the epigenetic regulatory mechanisms that promulgate the NPC phenotype. Here we obtained a comprehensive understanding of the 3D genome organization of nasopharyngeal cancer through Biop-C analysis, which revealed complex 3D genome organization patterns at oncogenes important in NPC.

## Results and Discussion

### A Genome-wide Map of 3D Genome Organization in Nasopharyngeal Cancer

We applied “Biop-C” to analyze three NPC tissue samples, *i.e.*, “S009”, “S010”, and “S012”. The tumor cores were collected by needle biopsies and immediately flash-frozen in liquid nitrogen (Fig. S1A). The tumor cores were approximately weighed in the range 3-10 mg, and the clinical characteristics of the samples are listed in Table S1. To prepare the tissue for analysis of 3D genome organization, we used a liquid nitrogen-cooled mini mortar and pestle for the pulverization of tissue within a microtube. This approach kept the biopsy sample frozen and reduced the risk of sample degradation. Additionally, this approach also provided the flexibility of performing the workflow from sample acquisition to HiC library preparation in a single microtube, which further minimized potential sample losses.

After pulverization, we processed the samples with a commercially available Arima *in situ* Hi-C kit. Briefly, the chromatin was fixed with formaldehyde in the nucleus and digested with a restriction enzyme. Then overhangs were filled in with biotinylated nucleotides followed by proximity ligation. After ligation, crosslinks were reversed, and the DNA was purified from protein. Further, the purified DNA was sheared to ~350 bp mean fragment size. Finally, the sequencing libraries were generated using low input swift bioscience Illumina-compatible adapters (Fig. 1A). The usage of the mini mortar and pestle followed by Hi-C is the key innovation of the Biop-C method. While this improvement in sample processing is a small change, it is highly effective in generating high-quality chromatin interaction libraries from needle biopsy clinical samples (Fig. S1B).

**Fig 1:**
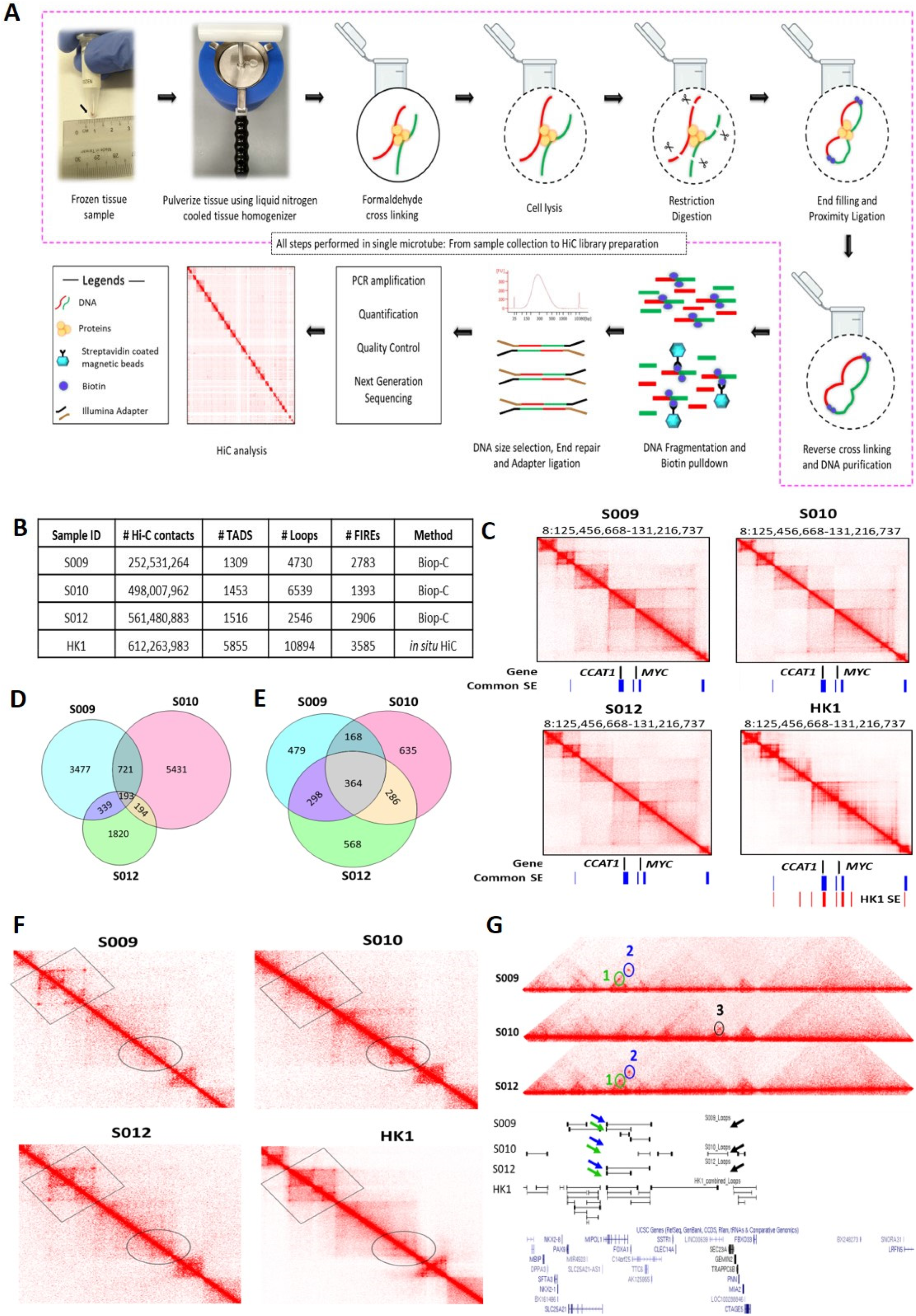
Biop-C method enables the study of high-resolution 3D Genome organization and chromatin interactome in solid tumor samples (A) Schematic overview of the Biop-C method. The frozen needle biopsy tissue is pulverized in a liquid nitrogen-cooled mini mortar, and then chromatin was fixed with formaldehyde in the nucleus. The samples were processed with commercial Arima genomics HiC Kit. Fixed cells were permeabilized using a lysis buffer supplied in Arima HiC kit and then digested with a restriction enzyme. Following restriction digestion, the sites were filled in with biotinylated nucleotides. The resulting DNA blunt ends were subsequently ligated. After ligation, crosslinks were reversed to remove proteins from DNA. Hi-C material was then sonicated using a Covaris Focused-Ultrasonicator M220 instrument to achieve fragment sizes of 300-500 bp. The sonicated DNA was double-size selected using Ampure XP beads, and the sequencing libraries are generated using low input Swift Bioscience Illumina-compatible adapters (B) Table of statistics of Biop-C and Hi-C data. #Hi-C contacts indicates number of mapped/valid junction reads of each library. #TAD indicates the number of TADs called from Biop-C and Hi-C data at 10kb resolution. #Loops indicates the total number of loops called from Biop-C and Hi-C data at 5kb, 10 kb, and 25 kb resolution and then merged. # FiREs indicates the number of FIREs called from Biop-C and Hi-C data at 10kb resolution. (C) Biop-C and Hi-C heat map showing 3D genome organisation at *CCAT1* and *MYC* gene. Genes are indicated in black color. The “common superenhancers”, shown in blue color, indicate the superenhancers present in all three NPC cell lines-HK1, C66-1, and HNE1 cell lines. The superenhancers present in HK1 cell lines are indicated in red color. The superenhancer datasets are obtained from Ke et al. 2017 [43] (D) Venn diagram showing the overlap of TAD boundaries between the patients inferred using the Arrowhead algorithm [9] at 10kb resolution. (E) Venn diagram depicting the overlap of Loops between the patients. The loops were called using the HiCCUPs algorithm [9] at 5kb, 10Kb, and 25 Kb resolution. The loops of the different resolutions were merged for this analysis. (F) Zoomed view of Biop-C Juicebox-visualized heat map of S009, S010, and S012 showing patient heterogeneity in chromatin interaction at *FOXA1* and *MIPOL1* gene. (G) UCSC genome browser [44] screenshots of genomic coordinates chr14:36,459,966-42,220,035. Loops 1 and 2 are present in S009 & S012, while loop 3 is present only in S010. Loop 1, Loop2, and Loop3 are represented in green, blue, and black colors, respectively.

Finally, we sequenced Biop-C libraries deeply by Illumina next-generation sequencing using a HiSeq4000 machine. Each library contained between 450 M and 922 M contacts (Table S2). We obtained more than 200 million mapped/valid junction reads (> 50% of total read pairs) for each library, reflecting that our Biop-C datasets are adequately complex [23] (Fig. 1B, Table S2). Furthermore, the low ratios of the number of *trans* to *cis* contacts indicate high library quality for all samples [24] (Table S2). Notably, in some samples (e.g., S009), only one lane of HiSeq4000 sequencing was sufficient to obtain a high quality Biop-C library.

We used Juicer for processing the resulting data, and the package Arrowhead was used to annotate Topologically-Associated Domains (TADs) genome-wide, while the package HiCCUPS was used to call loops [9]. Heatmaps were visualized with Juicebox [25]. We were able to successfully call TADs and loops from our libraries (Fig. 1B, Table S3 & S4). The TADs were clearly identifiable at a resolution of 10 kb. Patient S009 had 1309 TADs, while patient S010 had 1453 TADs and patient S012 had 1516 TADs (Fig. 1B). Loops could be identified at 5kb, 10 kb, and 25 kb resolutions in all datasets, and these loops were all merged together for subsequent analyses. Patient S009 had 4730 merged loops, while patient S010 had 6539 merged loops, and patient S012 had 2546 merged loops.

Moreover, because Frequently Interacting Regions (“FIREs”) are a new type of chromatin interaction landmark associated with super enhancers and tissue specific chromatin interactions [40], we used FIREcaller R package [45] to call Frequently interacting regions (FIREs). We identified 2783 FIREs in NPC sample S009, 1393 FIREs in sample S010 and 2906 FIREs in sample S012 from our Biop-C data. We also called FIREs from NPC cell line HK1 and identified 3585 FIREs (Fig. 1B).

Next, we examined the chromatin interactions around important oncogenes in NPC, such as MYC [26] and epidermal growth factor receptor (EGFR)[27–29]. The c-Myc is overexpressed in 76% of the NPC patients. The patients with c-Myc positive tumors had a longer disease-free period [26]. EGFR is highly expressed in nearly 85 % of NPC patients and associated with a significantly poorer prognosis in patients with advanced nasopharyngeal cancer than in patients without EGFR overexpression [28,29]. We found that EGFR is marked by three super enhancers in HK1 cell line, two of these three SE are localized upstream, i.e., near the start transcription site, and the third one is located in an intron (Fig. S1D).

Finally, for comparison with a typical Hi-C library, we generated two replicates of HK1 NPC cell line by traditional *in situ* Hi-C (Arima kit, [9,30]). Visual inspection of coverage normalized Biop-C heat maps of three NPC tissue samples, and Hi-C maps of HK1 cell line showed that the libraries were largely similar to each other, although we also noted that certain loci contained differences suggesting patient heterogeneity which we explore further in the next section of this manuscript (Fig 1C,1F,1G, S1C, S1I-J). Overall, our successful detection of TADs, loops and FIRES suggest that our Biop-C method can generate high-quality genome-wide chromatin conformation maps from the solid tumor needle biopsy samples.

### Patient heterogeneity in chromatin interactions

The question of patient heterogeneity is relatively unexplored in chromatin interaction analyses. In our previous research investigating chromatin interactions at the *TP53* and *MYC* loci, we observed that some chromatin interactions at MYC and TP53 could be detected in bone marrow and peripheral blood samples but not all chromatin interactions that are observed in K562 cells were detected in clinical samples [31].

To investigate potential patient heterogeneity, we compared the TADs and Loops between the patients’ Biop-C heatmaps using the Jaccard Index. We calculated Jaccard’s similarity coefficient (Jaccard Index, JI) to measure the overlap between the called TADs and loops in three Biop-C matrices. The resulting JI value indicates the fraction of shared TAD boundaries and loops between the patients. We observed that 38-42% of the TADs and 53-64 % of loops are shared in the three patients (Fig. S1L).

Next, we examined individual specific chromatin interactions. We found the chromatin interactions for the genomic locations *EGFR, PTPN1, DDIT4, MIR205HG, PDGFA, MALAT1, CAV2, NOTCH1, TEAD1, TP63, RUNX1*, *CCAT1, MYC,* and *YAP1* (Fig. S1C) are similar and *FOXA1, MIPOL1, SP4, SGCZ, MROH9, FMO1, FMO2, FMO1, FMO4,* and *FMO6P* gene are different (Fig. S1I, J). In one example, we observed that two loops i.e. loop 1 and loop 2 are present near *FOXA1* and *MIPOL1* genes in S009 and S012, which are thought to be tumor suppressors in nasopharyngeal cancer. However, these loops were absent in S010 (Fig 1F, G). In another example we observed a loop only in S009 and absent in S010 and S012 near miR383, which is considered an excellent diagnostic biomarker for head and neck cancer [35] (Fig. S1J).

Next, as we had observed patient-specific chromatin interactions, we investigated the tissue-specificity of these chromatin interactions. Thus, we characterized the similarities and the differences between the NPC landscape and other tissue types. We compared the Biop-C heatmaps and the Hi-C heatmap from HK1 with the previously published Hi-C heatmaps in human cell lines such as K562 (chronic myelogenous leukemia cell line), HAP1 (near-haploid cell line), IMR90 (fetal lung fibroblast cell line), KBM7 (chronic myelogenous leukemia), HUVEC (human umbilical vein endothelial cell line), RPE1 (retinal pigment epithelium cell line), GM12878 (lymphoblastoid cell line), NHEK (normal human epidermal keratinocytes), HeLa (human cervical carcinoma cell line), HCT116 (colon cancer cell line), HMEC (mammary epithelial cell line at a representative four genomic regions (Fig. S1E-H).

We observed that the HiC heatmaps of K562, HAP1, IMR90, KBM7, HUVEC, and RPE1 show a similar pattern as the Biop-C heat maps of S009, S010, S012, and HK1 Hi-C heatmap for the genomic locations around *MYC* and *CCAT1*, but differences in the profile were observed in GM12878, NHEK, HeLa, HCT116, and HMEC (Fig. S1E). The genome organization at *RUNX1* locus was similar to our patient’s Biop-C heatmaps in K562, HAP1, IMR90, RPE1, NHEK1, HeLa, HCT116, and HMEC but different in KBM7, HUVEC, and GM12878 (Fig. S1F). However, the genomic patterns for the *PTPN1* locus were found to be similar in all the Hi-C and patient’s Biop-C heat maps (Fig. S1G). On the other hand, the genomic patterns for the *MALAT1* locus did not show any similarity and were different in all the Hi-C, and Biop-C heat maps (Fig. S1H). We conclude that certain chromatin interactions in nasopharyngeal cancer are common across tissue types (*PTPN1*) and certain regions are tissue specific (*MALAT1*).

The observation of patient heterogeneity and tissue specificity in TADs appears to contradict earlier observations that TADs are primarily conserved across different human cell types and possibly even across different species [9,30,36]. However, Sauerwald analyzed 137 Hi-C samples from 9 studies and observed significant TAD variations across human cell and tissue types [37], suggesting that while there are common TADs and loops, there are also TADs and loops that vary across patient samples and tissue types.

### Super enhancers are associated with Frequently Interacting Regions and loop to genes

Super enhancers (SE) are regions of the DNA which enhances the transcription of target genes. These are comprised of group of enhancers which are at close proximity to each other and are marked by high enrichment of H3K27ac histone modification [38]. In previous research, we and others have shown that super enhancers can regulate distant genes via long-range chromatin interactions [31,39]. Moreover, frequently interacting regions (FIREs) have been reported to form at genomic regions also enriched by super enhancers [40]. Consequently, we wished to understand the relationship between super enhancers and chromatin interactions in nasopharyngeal cancer.

As the biopsy samples were too small for us to obtain both H3K27ac ChIP-Seq data as well as Biop-C data, we identified SE from NPC cell lines HK1, C666-1 and HNE1 using the ChIP-Seq data from [41] and we found that 298 SE were common in all the three cell lines (Table S5). We reasoned that these “common” super enhancers that are present in all cell lines examined will most likely also be present in the patient samples examined. We then used these common super enhancers and associated them with chromatin loops obtained from the Biop-C data of the patient samples as well as from the HiC data of HK1 cell line. As a result, we found that these SE are highly associated with chromatin loops. In sample S009, S010 and S012, the association of SE with chromatin loops were 54%, 57% and 41% respectively (Fig. 2A).

**Fig 2:**
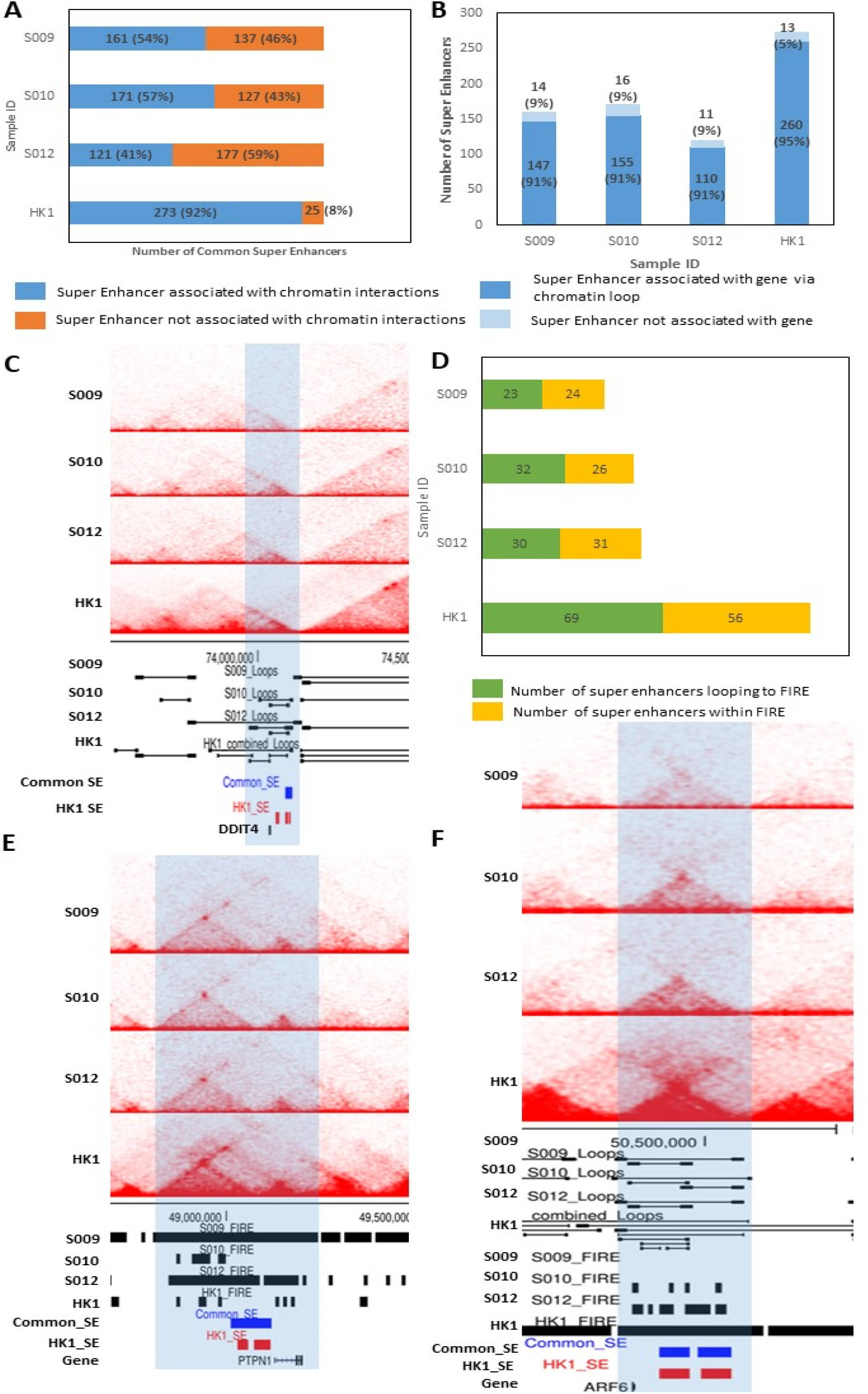
Association of super enhancers with chromatin interactions and genes. (A) Graph showing the number of common SE associated with chromatin interactions in NPC patient sample S009, S010, S012 and cell line HK1 (blue) and number of SE which are not associated with chromatin interactions (orange)(B) graph showing number of SE associated with distant genes via chromatin loops in NPC patient sample S009, S010, S012 and cell line HK1 (dark blue) and number of SE which do not link to distant genes via chromatin loops (light blue)(C) Biop-C heatmaps for NPC sample S009, S010, S012 and Hic heatmap for cell line HK1 for the *DDIT4* gene locus where a SE links to the gene through a chromatin loop (D) graph showing the number of SE associated with FIREs via chromatin loop (green) and SE which are within FIRE (yellow) (E) Biop-C heatmaps for NPC sample S009, S010, S012 and Hic heatmap for cell line HK1 showing a SE (blue: common SE, red: HK1 SE) and *PTPN1* gene within the same FIRE (F) Biop-C heatmaps for NPC sample S009, S010, S012 and Hic heatmap for cell line HK1 showing a SE looping to FIRE at the *ARF6* gene locus.

We further observed that more than 90% of these chromatin loops are associated with genes on the other side of the chromatin loop. In S009, 95% of SE associated chromatin loops are also linked to distant genes. In S010 as well as S012, 91% of SE associated loops are linked to genes (Fig. 2B). We also associated these SE to chromatin loops in one of the NPC cell line HK1 and found that 92% of common super enhancers are associated with chromatin loops in HK1 and 91% of these loops link SE to distant genes (Fig. 2A, B, Table S7). For example, *DDIT4* gene which is known to be over-expressed in NPC cell lines [41] is linked to a distant SE in patient sample S010, S012 and NPC cell line HK1 (Fig. 2C). We also repeated the above analysis with HK1 super enhancers and found that 85% of HK1 SE were associated with chromatin loops in HK1 cell line and 94% of these super enhancers were linked to distant genes via chromatin loops (Supplementary figure S2A, S2B). Next, we categorized SE which were at close proximity (less than 15kb from a gene) to a gene and were called “proximal” SE and the ones which are away from the gene and associated via chromatin loops were called distal SE. Out of 298 SE tested we found 27 proximal SE and 147 distal SE in S009, 155 distal SE in S010, 110 distal SE in S012 and 260 distal SE in HK1 (Supplementary table S7, S8). 48 genes were associated with proximal SE (Supplementary table S8) while 356 genes in S009, 421 genes in S010, 291 in S012 and 944 genes in HK1 were associated with distal SE (Supplementary table S7). There were also some genes that have both proximal and distal SE: In S009 we found 10 genes; in S010 as well as S012 we found 6 genes and in HK1 we found 18 such genes (Table S8). For example, *MACF1* gene in all the 3 NPC samples as well as HK1 cell line has distal as well as proximal SE (Table S8). We conclude that SE are highly associated with chromatin loops to distal genes in NPC.

Subsequently, we wanted to associate these SE with FIREs called (Table S6) from the Biop-C data from patient samples as well as HK1 HiC data. We could recognize 24 common SE in S009, 26 in S010, 31 in S012 and 56 in HK1 which are within a FIRE. Upon combining the chromatin loops data with FIRE calling, we found 23 SE in S009, 32 in sample S010, 30 in S012 and 69 SE in HK1 cell line that loops to a FIRE (Fig. 2D, Table S9). We also found SE which fall within the same FIRE as an oncogene like *PTPN1* which is also known to be over-expressed in NPC cell lines [41](Fig. 2E) as well as SE which loops to a distant FIRE containing *ARF6* gene whose over-expression can be correlated with metastasis and invasion in several cancers [42]. We conclude that SE are associated with FIREs in NPC.

Taken together, our new Biop-C method is suitable for interrogating needle biopsy patient samples, and more generally, situations of limited tumor sampling when surgical biopsies may be technically difficult. Using Biop-C, we examined chromatin interactions in 3 nasopharyngeal cancer patient samples, which allowed us to identify super enhancers associated with FIREs and which loop to important oncogenes. We also demonstrated patient heterogeneity in chromatin interactions in these patient samples, as well as tissue specificity. Hi-C libraries from an NPC cell line, HK1, showed differences compared with chromatin interactions in the patient samples. These differences could arise due to different subtype of NPC. Our results indicate the necessity of interrogating chromatin interactions in actual patient cancers, which Biop-C enables. Our results demonstrate the utility and importance of Biop-C as a method for understanding cancer 3D genome organization. In the future, we anticipate that the versatility of Biop-C will also allow us to interrogate perturbations of chromatin gene regulation in patients undergoing therapeutic interventions.

## Methods

Detailed methods are given in the Supplementary Materials. Briefly, the biopsy samples were obtained from NPC patients using an 18 gauge needle and flash frozen. The frozen samples were subjected to the Biop-C method for the generation of the Biop-C matrix. HK1 were cultured at 5% CO2 at 37°C in Roswell Park Memorial Institute (RPMI) 1640 media (Hyclone) supplemented with 10% heat-inactivated Fetal Bovine Serum (FBS; Hyclone) and 1% penicillin/streptomycin (Hyclone). The Hi-C heat maps of HK1 were prepared using Arima HiC kit and sequenced 150 bases paired-end on the Illumina HiSeq 4000. Hi-C data were aligned and processed by Juicer (version 1.5.7) [9]. The reference genome was hg19. The heat maps are visualized using Juicebox [25].

## Data deposition

We are in the process of submitting our data to GEO, dbgap and controlled short read archive.

## Data availability

All relevant data supporting the key findings of this study are available within the article and its Supplementary Information files or from the corresponding author on reasonable request.

## Acknowledgements

This research is supported by the National Research Foundation (NRF) Singapore through an NRF Fellowship awarded to M.J.F (NRF-NRFF2012-054) and NTU start-up funds awarded to M.J.F. This research is supported by the RNA Biology Center at the Cancer Science Institute of Singapore, NUS, as part of funding under the Singapore Ministry of Education Academic Research Fund Tier 3 awarded to Daniel Tenen (MOE2014-T3-1-006). This research is supported by the National Research Foundation Singapore and the Singapore Ministry of Education under its Research Centres of Excellence initiative.

## Author contributions

S.A. and M.J.F conceived the research idea. M.J.F, S.A., and R.C. contributed to the study design. S.A. and R.C. performed bioinformatics analysis of the Hi-C. S.A. performed manual curation of the Hi-C data. R.C performed FIRE calling and ChIP-Seq analysis on published data to identify S.E.s and their looping patterns in NPC. X.Y, J.T., W-Q.C and B.C.G. provided NPC. clinical samples. S.A., R.C, and M.J.F reviewed the data. S.A., R.C, B.C.G. and M.J.F wrote the manuscript. All authors reviewed and approved the manuscript.

